# Biosonar spatial resolution along the distance axis: revisiting the clutter interference zone

**DOI:** 10.1101/2020.02.27.967919

**Authors:** Peter A. Wagenhäuser, Lutz Wiegrebe, A. Leonie Baier

## Abstract

Unlike all other remote senses like vision or hearing, echolocation allows estimating the distance of an object. Not only have echolocating bats and toothed whales been shown to measure distance by echolocation extremely precisely, distance information is even topographically represented by a neuro-computational map in bats’ auditory cortex. This topographic representation and the corresponding tuning of cortical cells to object distance suggests the bats may be able to perceptually resolve multiple, simultaneously present objects along the distance axis. Here we use a novel psychophysical paradigm with complex phantom targets to quantity spatial resolution along the distance axis in the echolocating bat *Phyllostomus discolor*. We show that our bats can indeed perceptually resolve objects along the distance axis when they are separated by about 40 cm (around a reference distance of 108 cm) along the distance axis. These results are well comparable to earlier work on bats’ clutter interference zone (Simmons et al., 1988) and confirm those results with a more robust psychophysical paradigm.

**Summary statement:** Echolocating bats perceive absolute distance to objects by measuring the time delay between call and echo. In addition, they possess spatial resolution along the distance axis.

## Introduction

Distance is important: from an ecological perspective, the knowledge about an animal’s distance from either prey or predator is vital. However, distance is particularly difficult to assess: we get the most explicit information about the distance from an object when we can touch the object, but ecology would tell us that this is usually too late. Thus we need to assess distance with our remote senses, i.e. vision or audition.

Vision and audition possess fundamentally different sensory epithelia. The eye explicitly encodes the azimuth and elevation of a point in space on the retina, which allows for distance perception (or ranging) via binocular cues (Erkelens and van Ee, 1998; Howard, 1995; Howard, 2002; Qian, 1997; Rogers and Graham, 1979). On the contrary, the ear encodes sound as a function of frequency, not space, in the cochlea. The location of a sound source must therefore be computed in the brain. To localize a sound source in azimuth, differences of sound level and arrival times between the ears are exploited (Rayleigh, 1907). To localize a sound source in elevation, direction-dependent spectral interference patterns are exploited that are imprinted by the head and pinnae (the head-related transfer function (Blauert, 1997)). However, these computations do not provide cues about the distance of the sound source, i.e. ranging cues.

In passive hearing, i.e. when perceiving sounds that originate externally, distance estimation is only possible when the sound source is quite close (Kuwada et al., 2010; Kuwada et al., 2015) or very well known (Zahorik and Wightman, 2001). Absolute distance estimation is further facilitated in reverberant environments (Bekesy, 1938; Mershon, 1975), possibly because direct-to-reverberant ratio of sound levels serves as an explicit ranging cue (Bronkhorst and Houtgast, 1999).

In active hearing, i.e. echolocation, distance to an object can be directly perceived. Echolocation is an active auditory sense where animals emit ultrasonic sounds and listen to the echoes returning from objects around them. Thus, unlike any other sense, echolocation allows measuring the time between sound emission and echo reception. This so-called echo delay explicitly encodes the distance to an object: with the speed of sound at 340 m/s, an echo delay of e.g. 5 ms corresponds to an object distance of 85 cm. Indeed early studies have shown how sensitive echolocating bats and toothed whales are for object range (Murchison, 1980; Simmons, 1971; Simmons, 1979).

In bats, the importance of distance, and its perceptual equivalent echo delay, is reflected in neural specialisations along the entire auditory pathway (Covey and Casseday, 1991; Covey and Casseday, 1999; Grothe et al., 1992): bats possess delay-tuned neurons that respond strongest when the bat receives echoes from an object at a specific distance. The delay tuning culminates in a topographic representation of echo delay in the cortex: there is a clear relationship between a neuron’s position inside the postero-dorsal auditory cortex and its preferred echo delay (Bartenstein et al., 2014; Hagemann et al., 2010; O’Neill and Suga, 1979). The existence of this neurally computed distance map suggests that bats may not only be able to accurately localize objects along the distance axis, but also may be able to resolve multiple (acoustically semi-transparent) objects separated only along the distance axis. In other words, can delay tuning be the sensory basis not just of range accuracy, but also of range resolution?

A related question was addressed by Simmons et al. (1988). The authors trained *Eptesicus fuscus* bats to detect an electronically generated phantom reflection in the presence of masking reflections from real ring-shaped objects at a similar distance. In a two-alternative, forced choice experiment (2AFC), bats were presented with the one masking object each to the left and right, but the phantom reflection was added only to one, randomly chosen side, which the bats had to approach in order to get a food reward. The target strength of the target reflection was adjusted to be about 11-18 dB above detection threshold without masking reflections. When the phantom target reflection had a (virtual) distance of e.g. 80 cm, bats’ performance dropped significantly when the masking rings were presented at distances between 50 and 90 cm (Simmons et al., 1988).

With this experimental paradigm Simmons et al. (1988) characterized a clutter interference zone along the distance axis, i.e., a range of distances where object detection is not independent of one another (clutter refers to non-target structures and echoes). The paradigm is unsuitable to quantify range resolution, however, because the bats only had to discriminate between a single reflection and two reflections (the side with only the ring vs. the side with both the phantom target and the ring). The bats could have used a range of different perceptual cues to solve the target-detection task: the rewarded side with two reflections provides (i) a higher overall target strength, (ii) a longer overall echo, and (iii) the interference between the target and masking reflection on the rewarded side will create a comb-filter spectrum (cf. Figure 5 of Simmons et al. (1988)).

**Fig. 1.**
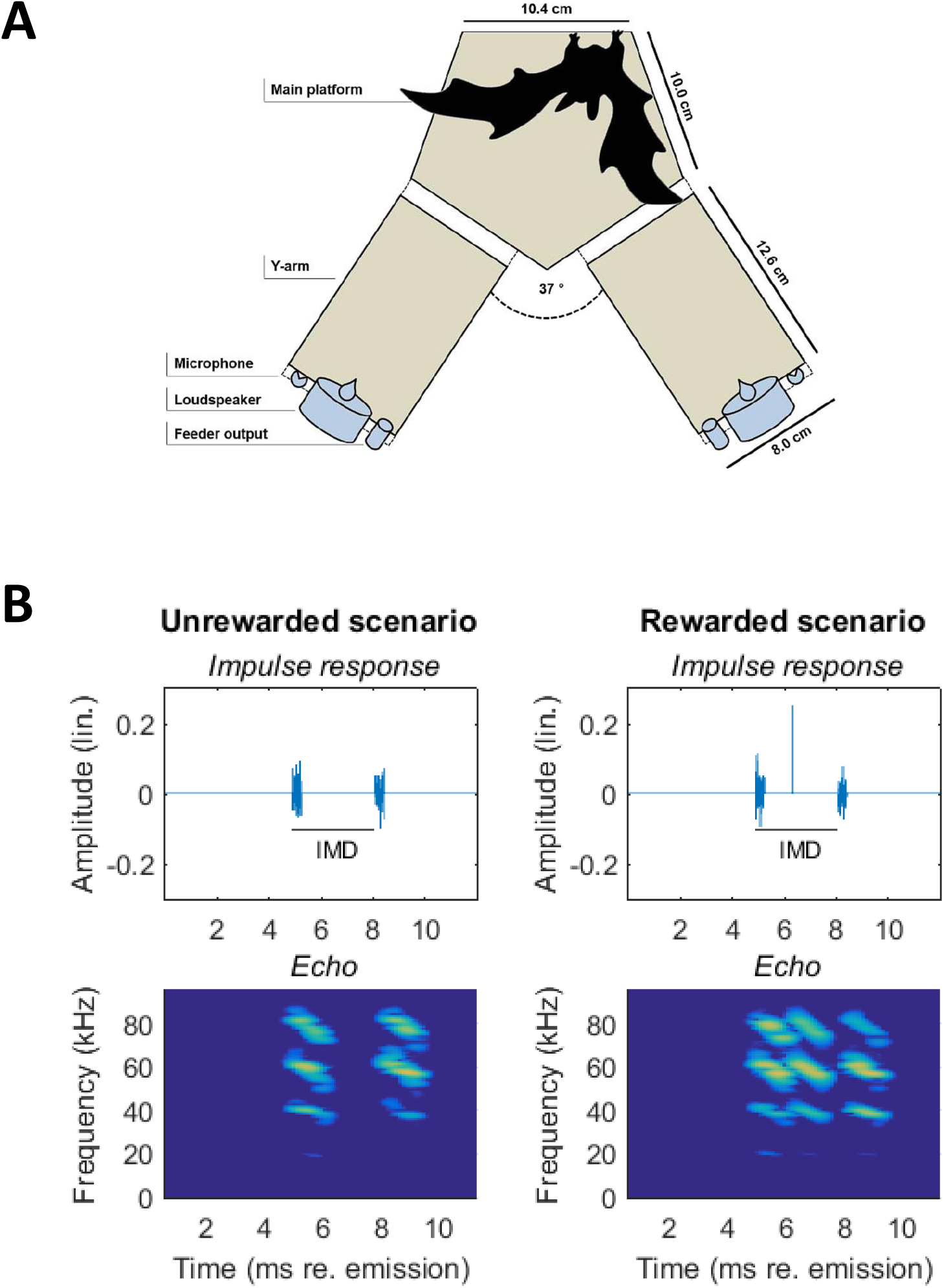
Setup and stimuli. **(A)** Auditory-virtual-reality setup: bats were trained to discriminate a virtual scenario without target from a virtual scenario with target. All bats learned to indicate the pseudorandomly chosen position of the rewarded scenario by crawling towards it from the main platform after echolocating towards both scenarios. Virtual scenarios were created by convolving recorded echolocation calls in real time with a pre-defined impulse response (see Methods). **(B)** Impulse responses (top) and echoes (bottom) of the virtual scenarios: the unrewarded scenario consists only of the two maskers (left); the rewarded scenario (right) contains the target reflection. The inter-masker delay (IMD) between the first and the second virtual masker was always geometrically centred around the target delay of 6.3 ms.

**Fig. 2.**
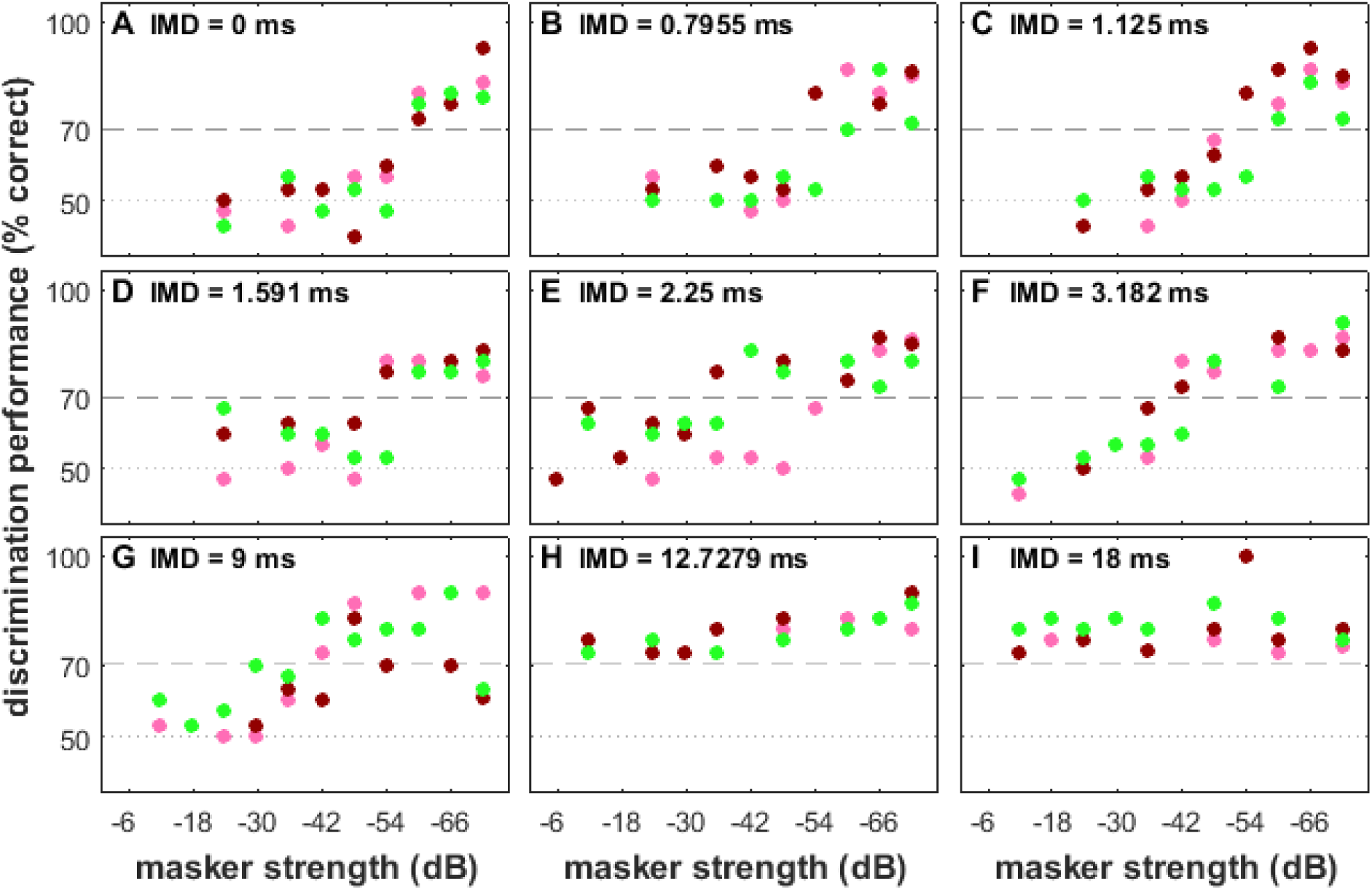
Psychometric functions of target detection performance at nine inter-masker delays (IMDs) Each coloured dot marks one bat’s discrimination performance across 30 trials (pink: bat 1, ruby: bat 2, green: bat3). Horizontal dashed lines at 50 and 70 % correct depict chance and significance level, respectively. For single bats, percent correct performance as a function of modulation depth was fitted with a sigmoidal function and the value of this fit at 70% was taken as threshold. **(A-G)** For IMDs up to 9ms, discrimination performance decreased with increasing masker strength, eventually dropping to chance level. **(H-I)** For IMDs higher than 9ms, discrimination performance did not drop below significance level throughout the range of tested masker strengths.

**Fig. 3.**
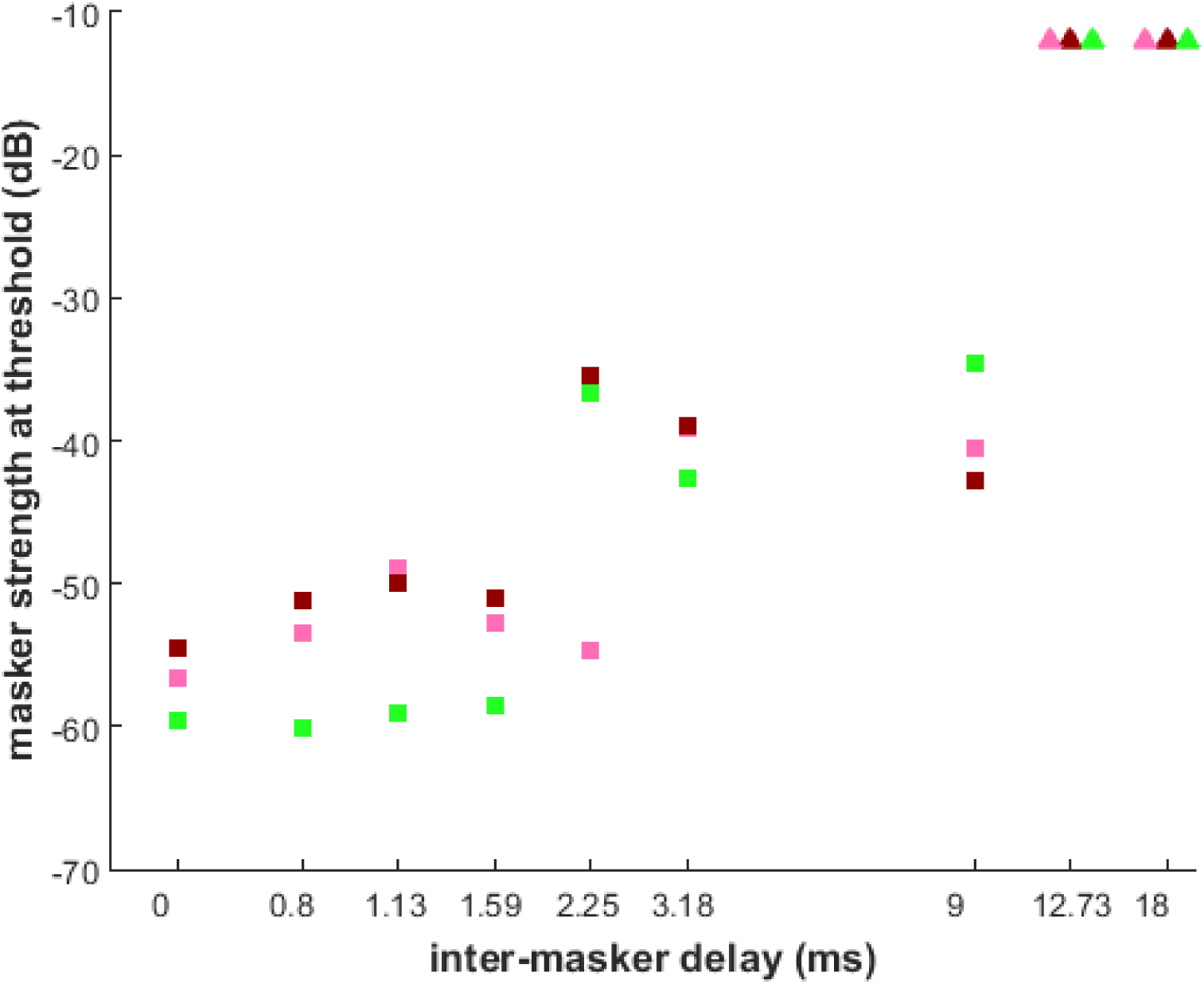
Masker strength at threshold as a function of inter-masker delay (IMD) Masking thresholds were extracted from psychometric functions in Fig.2. Each coloured square marks one bat’s masking threshold for IMDs between 0 ms and 9ms (pink: bat 1, ruby: bat 2, green: bat3). Coloured triangles indicate that masking thresholds at higher IMDs are at least −12 dB, because discrimination performance did not drop below significance level throughout the range of tested masker strengths (cf. Fig. 2H-I). Note the logarithmic ordinate.

**Fig. 4.**
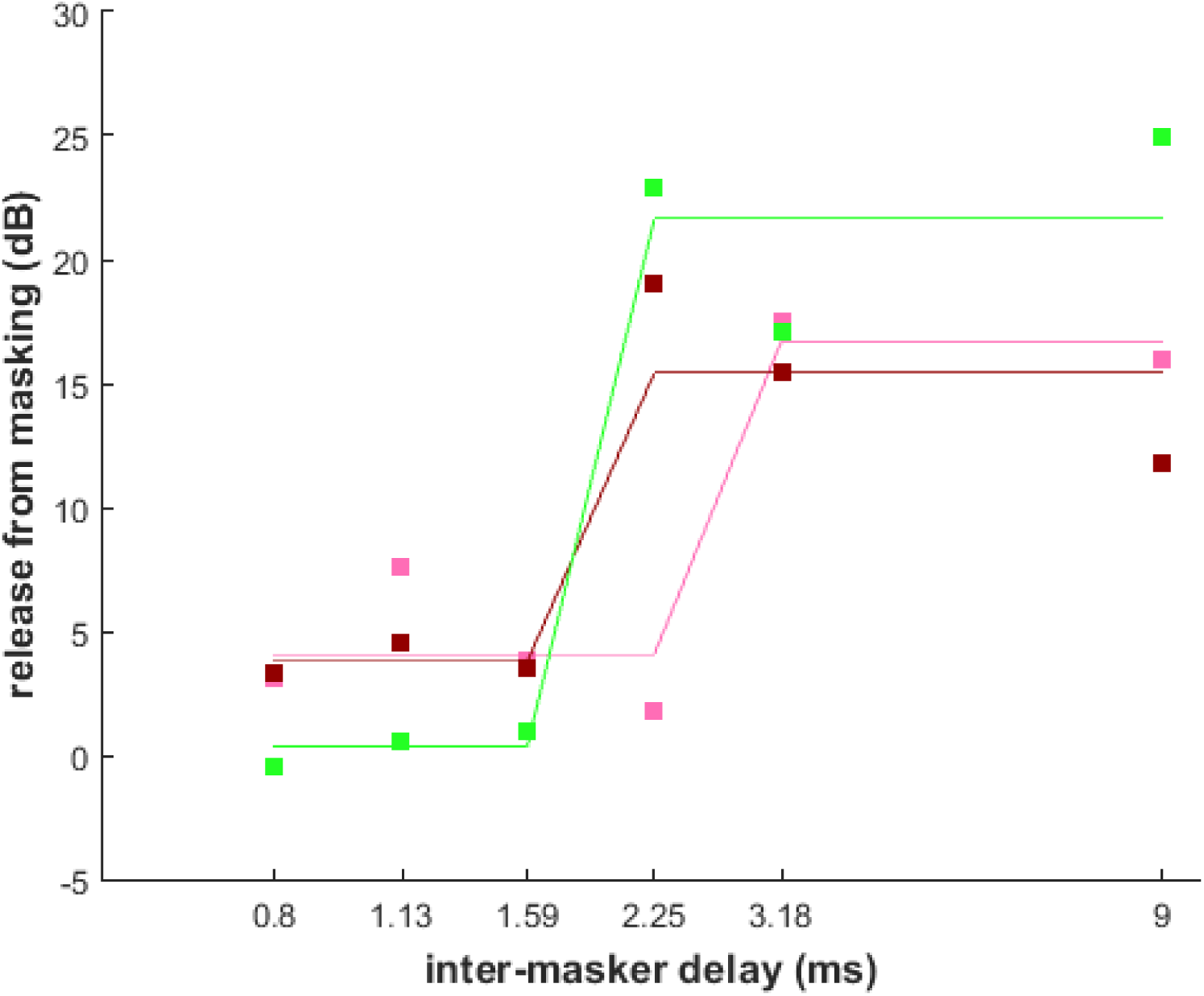
Release of masking as a function of inter-masker delay (IMD) Release-of-masking values were calculated from masking thresholds in Fig. 3. Each coloured square marks one bat’s release-from-masking value at the respective IMD (pink: bat 1, ruby: bat 2, green: bat3). Coloured lines are sigmoidal fits for single bats. The resolution limit for each bat is extracted as the turning point of the fit. Note the logarithmic ordinate.

**Fig. 5.**
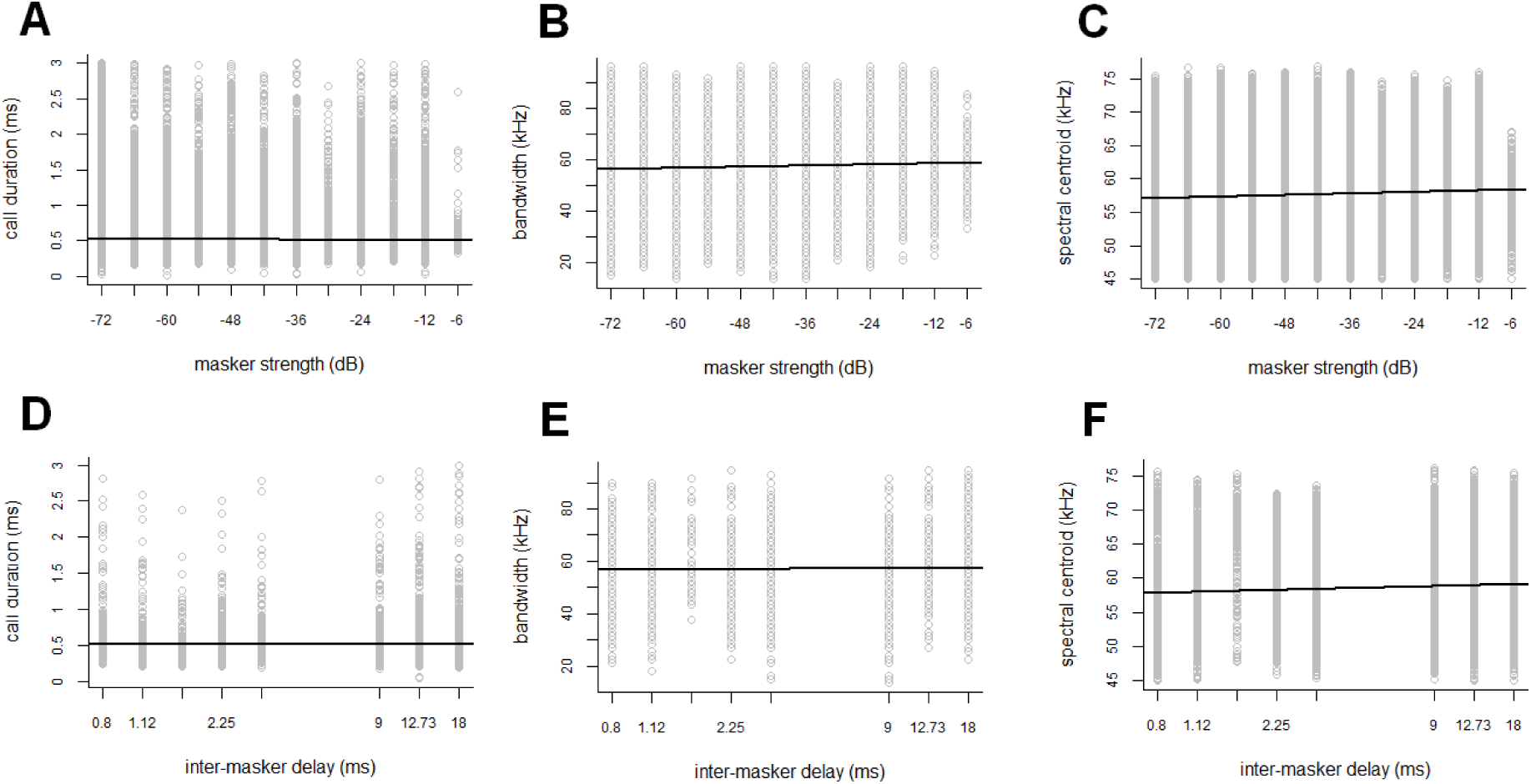
Linear regression analysis of temporal and spectral call parameters. Effects of masker strength and inter-masker delay (rows) on bats’ call parameters (columns). Analysis was carried out only for those trials that bats had solved correctly. For analysis of inter-masker delay-effects (bottom row), only calls from trials with masker strength close to the individual bat’s detection limit were used.

We aimed to design an experimental paradigm that precludes the above-mentioned perceptual cues that may complicate the interpretation of the results by Simmons et al. (1988) and thereby allows for the quantification of spatial resolution along the range axis. Lord Rayleigh provided a clear definition of spatial resolution: two closely spaced light sources are spatially resolved when there is a detectable dip in their joint light diffraction patterns (Rayleigh, 1879; Westheimer, 2005).

Here we used a virtual environment to quantify the range-resolution limit for bat biosonar, following Lord Rayleigh’s definition of spatial resolution. We generated patterns of reflections that allowed for detecting the perceptual dip between two objects that were adjoined along the distance axis. We report on our results from a psychophysical detection experiment with echolocating *Phyllostomus discolor* bats. We demonstrate that with a delay difference of ∼2.2 ms at a reference delay of 6.3 ms, the bats can listen into the dip between two masking reflections and detect a probe reflection. We conclude that for a target distance of 1 m, *P. discolor* bats have a range-resolution limit of about 37 cm.

## Materials and Methods

### Animals and permit

We used three adult male individuals of the neotropical omnivorous bat species *Phyllostomus discolor*. These bats emit short (<3 ms), downward frequency-modulated (FM), multi-harmonic echolocation calls covering the frequency range between 45 and 100 kHz (Rother and Schmidt 1982). Bats were kept at the bat facilities in the Department Biology II of the Ludwig-Maximilians-University in Munich (12 h night / 12 h day cycle, 65-75% relative humidity, 28°C) with unlimited access to water at all times. On free days, the bats had ad libitum access to mixed fruit and mealworms (larval form of Tenebrio molitor) supplemented with oat, safflower oil, baby formula, minerals and vitamins (Vitakalk®). During training periods, the bats were with fed a pulp from fruit and supplementals in the experiment. All experiments complied with the principles of laboratory animal care and were conducted under the regulations of the current version of the German Law on Animal Protection (approval 55.2-1-54-2532-34-2015, Regierung von Oberbayern).

### Experimental setup

The experiments were performed on an open Y-maze inside a dark, echo-attenuated chamber. The 3D-printed Y-maze (see Fig. 1A) consisted of a pentagram-shaped starting area (side length 10 cm) and two arms (width and length 8×12.6 cm) and was covered in removable cloth. The loudspeakers and microphones as well as the food dispensers were mounted at the end of each arm. The experimenter was outside the chamber and observed the experiment via an infrared camera (Abus® TV6819) and headphones emitting heterodyned versions of the microphone signals. Stimulus presentation and data recording were controlled via a custom MatLab® R2007b application (The Mathworks, Inc., Natick, MA) and soundmexpro.

### Virtual scenario generation

Bats were trained to detect a virtual target flanked by two virtual maskers. All target and masker reflections were implemented as virtual reflections, generated by a real-time stereo convolution engine that calculated complex echoes from the bats’ ultrasonic emissions. The structure of the impulse responses (IRs) loaded into the convolution engine defined the echo-acoustic properties of the virtual reflections.

The unrewarded IR (Fig. 1B, left) consisted of two virtual maskers alone; the rewarded IR consisted of two virtual maskers surrounding a virtual target reflection (Fig. 1B, right). The maskers were implemented as short (∼300 µs) noise bursts; the target was implemented as a simple reflector. For each trial in the psychophysical procedure, the noise bursts were refreshed. This ensured that there was no systematic spectral interference between the masker- and target reflections which would have generated unwanted spectral cues. Echoes as they are generated with these complex IRs excited by a standard *P. discolor* echolocation call are shown in the bottom panels of Fig. 1B. We generated the scenarios by convolving a call recorded through the microphones with either of two IRs (maskers without and with target) and playing back the resulting virtual echo via the loudspeakers. Every change a bat chose to make in its emission sequence (e.g. change in call timing, call spectrum, or call direction) was immediately reflected in the echoes.

Specifically, the bat’s ultrasonic emissions were picked up by two microphones (SPU0410LR5H-QB, Knowles Corporation, Itasca, IL, USA) mounted 45° left and right relative to the bat’s starting position on the Y maze. The microphone signals were amplified (octopre LE, Focusrite plc, Bucks, United Kingdom) and fed into the inputs of two real-time digital signal processors (RX6, Tucker Davis Technologies, Gainesville, FL, 260 kHz sampling rate). In one processor, the signal was convolved with the rewarded IR (containing both the masker- and the target reflections) while in the other processor, the signal was convolved with the unrewarded IR (containing only the masker reflections). A constant base delay preceded the IRs in both processors such that the overall delay of the target reflection, including digital delays and acoustic delays from the bats’ emissions travelling to the microphones and the echoes travelling from the loudspeakers back to the bats amounted to 6.3 ms. The delay between the first and the second virtual masker (inter-masker delay IMD) was set by the experimenter. The masker delays were always geometrically centred around the target delay of 6.3 ms. This was done because the sharpness of cortical tuning to echo delay appears to scale with absolute echo delay (Greiter and Firzlaff, 2017; Hagemann et al., 2010; Suzuki and Suga, 2017). The outputs of the real-time processors were connected via a stereo amplifier (Harman Kardon HK 6150; Harman Deutschland, Heilbronn, Germany) to two ultrasonic speakers (Peerless XT25SC40-04, Tymphany HK Limited, San Rafael, USA). The target strength of the target reflection was fixed at −12 dB; the root-mean-square target strengths of the maskers were varied between −72 dB and −12 dB to obtain a psychometric function and a threshold for that masker strength where the signal was just detectable. Psychometric functions and thresholds were acquired for IMDs of 0 ms (reference condition), 0.80 ms, 1.13 ms, 1.59 ms, 2.25 ms, 3.18 ms, 9.00 ms, 12.73 ms, and 18.00 ms and for each of three bats (see below). The IMD is the onset difference between the first and second masker reflection.

### Behavioural procedure

Training/recording sessions (one to three per day) each lasted ten minutes. Bats were trained on five days per week, followed by a two-day break. The experiment followed a two-alternative, forced-choice paradigm (2AFC) with food reinforcement. Once a bat sat in the starting area of the Y-maze, presentation of the IRs was switched on. The position of the target reflection (left or right) was pseudorandom from trial to trial. Bats had to echolocate to identify and move towards the IR that contained the target reflection, where they were rewarded as soon as they reached the corresponding feeder. Once a bat had learned this task with very faint maskers (−72 dB and >70% correct choices on 5 consecutive days), the strength of all four masking reflections was increased, making the detection task more difficult. Starting each session with three consecutive trials presenting the weakest maskers (−72 dB), data acquisition proceeded by increasing the masker strengths in steps of 6 dB until the bats could not detect the target at all, and then restarting at very low masker strengths until the daily sessions were completed. Testing for one IMD set was completed when at least 30 trials were obtained per masker strength and bat.

### Behavioural data analysis

Percent correct performance of the animals as a function of masker strength were fitted with a sigmoidal function and the value of this fit at 70% was taken as threshold (for p<0.05 in a binomial test cf. Fig. 2). The threshold for a specific IMD is the masker strength that just allows a bat to reliably detect the target in the presence of the maskers. For each bat, we calculated release-from-masking values for IMDs between 0.7955 ms and 9ms as the difference between the respective threshold and the reference threshold at an IMD of 0 ms. Release-from-masking values were fitted with a sigmoidal function whose turning point determined the resolution limit of each bat.

### Acoustic analyses

The echo properties of each scenario depended both on the properties of the IRs and critically on the properties of the echolocation calls that the bats emitted. Therefore we verified the echolocation-call properties with acoustic analyses performed with custom MatLab® R2015a programs. We performed further statistical analysis with the freeware R version 3.5.1 (R Core Team, 2018).

During the psychophysical experiment, the recorded call sequences were saved in a 3-s stereo ring buffer (192 kHz sampling rate, 24-bit resolution; Fireface) parallel to the virtual-scenario production. Offline, we band-pass filtered the stereo recordings at 100 Hz to 20 kHz applying a 2nd order butterworth filter. We applied a synthesized echolocation call (multiharmonic FM-downward sweep of 1 ms duration with a fundamental frequency ranging from 21–18 kHz) as a matched filter to separate echolocation calls from other transient events like click sounds made by the setup when the bats stepped onto one of the response platforms. Temporal and spectral call parameters were taken from the channel corresponding to the rewarded scenario. We calculated the −10 dB call duration. We calculated the −20dB bandwidth from minimum and maximum frequencies. We calculated the spectral centroid (weighted mean of frequencies present in the signal) from a time-averaged spectrogram with a 1500 Hz binwidth.

We performed statistical analysis with the freeware R, version 3.5.1 (R Core Team, 2018). First, we investigated the relation between a given call parameter (y) and masker strength (x). We only analysed calls recorded during those trials that the bats had solved correctly. Second, we investigated the relation between a given call parameter (y) and inter-masker delay (x). Here, we again analysed only calls recorded during trials that the bats solved correctly, and only during those trials where the maskers had been presented at an individual bat’s detection limit, i.e. a target strength that just allowed for target detection. We fitted linear mixed effects models by the maximum likelihood method with a Gaussian error distribution using the *lmer* function of the R-package *lme4* (Bates et al., 2014). As response variable we used different call parameters. The respective response variable was first modelled as a function of the fixed effect masker strength (z-transformed, so that its mean was 0 and its standard deviation was 1), and second as a function of the fixed effect inter-masker delay (also z-transformed and log-transformed). We included *session ID* as random factor to account for repeated measures within the same sessions, and *bat* as second random factor to account for individual differences. In the first model, the effect of masker strength was grouped by IMD, because we used call data from all tested IMDs. We verified model assumptions inspecting quantile-quantile plots of the model residuals. We used improper prior distributions, namely p(β) ∝ 1 for the coefficients, and 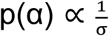 for the variance parameters. To obtain the posterior distribution we simulated 1000 values from the joint posterior distribution of the model parameters with the *sim* function of the R-package *arm* (Gelman and Hill, 2006). We first extracted 95% credible intervals (CrI) around the regression line by calculating 1000 fitted values for the respective range of x-values. We then investigated the posterior predictive distribution in order to report where future observations would scatter around the regression line. For each simulated fitted value, we simulated one new y-value and extracted the 95% interval of the simulated predicted distribution.

## Results

### Behavioral response

Three male bats (*Phyllostomus discolor*) learned to discriminate between a virtual scenario consisting of two masker reflections and a virtual scenario consisting of two masker reflections plus the target reflection. We used the behavioural response of the bats to assess the masking thresholds, i.e. the highest masker strength that still let the bats detect the target reflection. For inter-masker delays (IMDs) ranging from 0 ms to 9 ms, the results of all bats confirmed our expectations for a psychometric function: discrimination was good at low masker strengths and deteriorated with increasing masker strength (Fig. 2A-G). For IMDs of 12 ms and 18 ms, however, discrimination performance remained above chance level regardless of masker strength (Fig. 2H-I). Consequently, for every bat we extracted one masking threshold per IMD from the psychometric functions for IMDs between 0 ms and 9 ms.

For IMDs between 0 ms and 9 ms, all bats reliably (70–90% correct choices; Fig. 2A-G) detected the target reflection when the maskers were very faint (masker strength of −60 dB and lower). In contrast, when the maskers were very loud (masker strength of −24dB and higher) none of the bats could solve the detection task for these IMDs (43–65% correct choices; Fig. 2A-G).

For IMDs between 0 ms and 9 ms, detection performance as a function of masker strength systematically changed with IMD. Masking thresholds remain around −50 to −60 dB for IMDs shorter than about 3 ms but improve rapidly when the IMD is increased further (Fig. 3).

### Distance resolution threshold

In order to derive a distance resolution threshold from the behaviourally obtained masking thresholds, we assessed the release from masking provided by the separation of the maskers. As outlined in the introduction, the two masker reflections are perceptually resolved in distance when there is a significant dip in their perceptual representation. This dip is probed with the target reflection. The dip is significant when the masking effect elicited by both maskers is significantly less than that elicited by one of the maskers. In the reference condition with an IMD of 0 ms, both maskers are presented simultaneously, acting as one masker. However, the noise power of both masker reflections adds up, making this one masking reflection 3 dB stronger. Consequently, the maskers are spatially resolved when the release from masking (relative to the IMD of 0 ms) is at least 3 dB. We calculated release-from-masking values as the difference between each respective masking threshold for IMDs between 0.8 ms and 9 ms and the masking threshold for an IMD of 0 ms. We extracted exact values as the turning points of fitted sigmoidal functions. On average, a release from masking larger than 3 dB is seen when the IMD exceeds about 2.2 ms (Fig. 4). Converting echo delay into distance measures, bats showed a distance-resolution limit of about 37 cm for a target distance of 1.07 m (6.3 ms reference delay).

### Acoustic analysis

The bats’ auditory percept depended not only on the echoacoustic features of the virtual scenarios themselves, but critically on how the bats ensonified them. We performed acoustic analyses of the echolocation calls used by the bats during the behavioural experiment to better understand which strategies the bats employed to solve the task.

First we tested whether fundamental temporal and spectral call parameters changed systematically when the task became more difficult for the bats, i.e., when the masker strength increased. We fitted a linear regression for three call parameters: call duration, bandwidth and spectral centroid (weighted frequency mean). The relationship between call parameters and masker strength is very poor (Fig 5 A-C).

Bat 1 used a mean call duration of 0.46 ms (SD 0.23 ms, n=20490) for a masker strength of −72 dB re. target and a mean call duration of 0.42 ms (SD 0.22 ms, n=3894) for a masker strength of −12 dB re. target. Bat 2 used a mean call duration of 0.64 ms (SD 0.28 ms, n=36980) for a masker strength of −72 dB re. target and a mean call duration of 0.60 ms (SD 0.28 ms, n=2453) for a masker strength of −12 dB re. target. Bat 3 used a mean call duration of 0.59 ms (SD 0.30 ms, n=18438) for a masker strength of −72 dB re. target and a mean call duration of 0.65 ms (SD 0.39 ms, n=2599) for a masker strength of −12 dB re. target.

Bat 1 used a mean call bandwidth of 57.23 kHz (SD 7.98 kHz, n=20490) for a masker strength of −72 dB re. target and a mean call bandwidth of 57.50 kHz (SD 7.84 kHz, n=3894) for a masker strength of −12 dB re. target. Bat 2 used a mean call bandwidth of 59.90 kHz (SD 13.27 kHz, n=36980) for a masker strength of −72 dB re. target and a mean call bandwidth of 61.01 kHz (SD 11.3 kHz, n=2453) for a masker strength of −12 dB re. target. Bat 3 used a mean call bandwidth of 54.42 kHz (SD 9.76 kHz, n=18438) for a masker strength of −72 dB re. target and a mean call bandwidth of 57.87 kHz (SD 10.13 kHz, n=2599) for a masker strength of −12 dB re. target.

Bat 1 used a mean spectral centroid (SC) of 63.26 kHz (SD 5.48 kHz, n=20490) for a masker strength of −72 dB re. target and a mean SC of 64.36 kHz (SD 5.52 kHz, n=3894) for a masker strength of −12 dB re. target. Bat 2 used a mean SC of 54.42 kHz (SD 5.00 kHz, n=36980) for a masker strength of −72 dB re. target and a mean SC of 55.65 kHz (SD 5.63 kHz, n=2453) for a masker strength of −12 dB re. target. Bat 3 used a mean SC of 51.84 kHz (SD 3.87 kHz, n=18438) for a masker strength of −72 dB re. target and a mean SC of 52.73 kHz (SD 4.44 kHz, n=2599) for a masker strength of −12 dB re. target.

Second, we determined whether bats changed their temporal and spectral call parameters systematically with inter-masker delay (IMD), i.e. below or above resolution limit. Again, we fitted a linear regression for three call parameters, but we used only data from those trials where masker strength was close to the masking threshold for the specific IMD and bat. As for masker strength, the relationship between call parameters and IMD is very poor (Fig 5D-F).

At masking threshold, bat 1 used a mean call duration of 0.42 ms (SD 0.16 ms, n=646) for an IMD of 0.8 ms (unresolved), a mean call duration of 0.42 ms (SD 0.15 ms, n=699) for an IMD of 3.2 ms (just resolved), and a mean call duration of 0.42 ms (SD 0.20 ms, n=715) for an IMD of 18.0 ms (clearly resolved). At masking threshold, bat 2 used a mean call duration of 0.65 ms (SD 0.35 ms, n=1083) for an IMD of 0.8 ms, a mean call duration of 0.59 ms (SD 0.21 ms, n=629) for an IMD of 3.2 ms, and a mean call duration of 0.63 ms (SD 0.31 ms, n=1449) for an IMD of 18.0 ms. At masking threshold, bat 3 used a mean call duration of 0.61 ms (SD 0.19 ms, n=223) for an IMD of 0.8 ms, a mean call duration of 0.55 ms (SD 0.23 ms, n=471) for an IMD of 3.2 ms, and a mean call duration of 0.59 ms (SD 0.34 ms, n=518) for an IMD of 18.0 ms.

At masking threshold, bat 1 used a mean call bandwidth of 56.45 kHz (SD 4.13 kHz, n=646) for an IMD of 0.8 ms (unresolved), a mean bandwidth of 57.78 kHz (SD 8.40 kHz, n=699) for an IMD of 3.2 ms (just resolved), and a mean bandwidth of 57.70 kHz (SD 8.74 kHz, n=715) for an IMD of 18.0 ms (clearly resolved). At masking threshold, bat 2 used a mean bandwidth of 55.68 kHz (SD 13.04 kHz, n=1083) for an IMD of 0.8 ms, a mean bandwidth of 58.19 kHz (SD 13.32 kHz, n=629) for an IMD of 3.2 ms, and a mean bandwidth of 60.65 kHz (SD 11.35 kHz, n=1449) for an IMD of 18.0 ms. At masking threshold, bat 3 used a mean bandwidth of 56.03 kHz (SD 12.60 kHz, n=223) for an IMD of 0.8 ms, a mean bandwidth of 55.78 kHz (SD 7.83 kHz, n=471) for an IMD of 3.2 ms, and a mean bandwidth of 57.05 kHz (SD 7.44 kHz, n=518) for an IMD of 18.0 ms.

At masking threshold, bat 1 used a mean spectral centroid (SC) of 68.20 kHz (SD 4.23 kHz, n=646) for an IMD of 0.8 ms (unresolved), a mean SC of 64.44 kHz (SD 5.91 kHz, n=699) for an IMD of 3.2 ms (just resolved), and a mean SC of 65.99 kHz (SD 4.93 kHz, n=715) for an IMD of 18.0 ms (clearly resolved). At masking threshold, bat 2 used a mean SC of 53.81kHz (SD 4.83 kHz, n=1083) for an IMD of 0.8 ms, a mean SC of 53.69 kHz (SD 4.95 kHz, n=629) for an IMD of 3.2 ms, and a mean SC of 55.02 kHz (SD 5.95 kHz, n=1449) for an IMD of 18.0 ms. At masking threshold, bat 3 used a mean SC of 51.40 kHz (SD 4.17 kHz, n=223) for an IMD of 0.8 ms, a mean SC of 50.19 kHz (SD 3.36 kHz, n=471) for an IMD of 3.2 ms, and a mean SC of 52.84 kHz (SD 3.14 kHz, n=518) for an IMD of 18.0 ms.

In conclusion, we found no evidence for an adjustment of call parameters; neither in response to masker strength nor in response to inter-masker delay (IMD).

## Discussion

Echolocating bats perceive absolute distance to objects by measuring the time delay between call and reflection. With the current psychophysical experiment we show that *Phyllostomus discolor* bats can also resolve multiple reflections along the distance axis. We used the target reflection as a probe to characterize the temporal auditory excitation pattern generated by the masking reflections. We show that the resolution limit is about 2.2 ms when the maskers are centred around a reference delay of 6.3 ms. This resolution limit is equivalent to a range of about 37 cm around a reference distance of 1.07 m.

In the following paragraphs we first discuss the ‘clutter interference zone’: (Simmons et al., 1988) in terms of experimental design and significance. Second, we consider the resolution limit in the context of previous experiments on object detection along the distance axis. Third, we examine the acoustic properties of the echolocation calls that the bats used throughout the current experiment and discuss their influence on performance.

### The clutter interference zone

As already mentioned in the introduction, the current experiments address similar issues as the experiments in the ‘clutter interference zone’: Simmons et al. (1988) trained bats to detect a virtual target reflection in the presence of masking reflections off a ring-shaped object, characterizing a range of distances where object detection is not independent of one another. However, with this paradigm it is unclear which strategy the bats may have used to solve the psychophysical task: besides perceptually resolving target- and masker reflections, the bats could also have evaluated a number of different perceptual cues (related to (i) overall target strength, (ii) overall echo duration, or (iii) spectral interference between the target and masker reflection, cf. Introduction). In criterion-free psychophysical procedures such as the 2AFC procedure (Green and Swets, 1966), the subject is free to choose the one perceptual cue (or combination of cues) that provides the highest success rate. By no means is it certain that the perceptual cues that the experimenter expected to probe are identical to those perceptual cues that the subjects actually use to solve the task. Therefore, stimulus design is critical.

The current experiment (cf. Fig. 1) was designed to preclude the use of additional perceptual cues: (i) the overall target strength for the unrewarded and the rewarded impulse response (IR) differed by the target strength of the target reflection, which was set to −12 dB, far below *P. discolor*’s threshold for amplitude discrimination (Heinrich et al., 2011); (ii) the overall duration of the stimulus was set by the inter-masker delay (IMD) and was the same for the unrewarded and the rewarded IR; (iii) the IRs of the masker reflections consisted of noise bursts that were repeatedly refreshed so that they did not create systematic spectral interference with the target reflection. We are confident that our current stimulus design prevented unwanted perceptual cues and let us actually probe biosonar resolution of objects along the distance axis.

Notably, however, the current paradigm requires much more measurements than the paradigm by Simmons et al. (1988). The clutter interference zone was determined with one psychometric function per bat (albeit at three different reference distances): the target reflection was set to a target strength just detectable by the bat without maskers and the bat’s performance was then measured as a function of masker position relative to the target. Here, we measured performance as a function of masker strength and recorded a complete psychometric function per bat for each IMD. The current results are therefore based on about ten times the number of data points per bat compared to the clutter interference zone experiment (cf. Fig. 2).

These crucial points notwithstanding, the resolution limit quantified here is quite similar to the limits of the clutter interference zone by Simmons et al. (1988). They found that for a target distance of 40, 80, or 160 cm, clutter interference zones for one bat extended about 25, 32, and 60 cm around the target distance, respectively (cf. Fig. 4 in Simmons et al. (1988)). We demonstrate here a resolution limit of around 37 cm at a reference distance of 1.07 m. The similarity of the results indicate that in the study by Simmons et al. (1988), bats may have relied on temporal-resolution cues to separate the target reflection from the clutter reflections, despite the presence of multiple other perceptual cues.

### Object detection along the distance axis

Aside from the direct comparison to formal psychophysical experiments, the current results belong in the context of object detection along the distance axis. Detecting a target in front of or behind a non-target is a common task in biosonar. To detect prey in clutter, i.e. among non-target structures, bats usually apply one of several foraging strategies (reviewed in Denzinger and Schnitzler (2013)). They either use other sensory systems such as vision or olfaction, they hunt only moving prey that generate peculiar echoes (flutter detection), or they eavesdrop on prey-generated sounds (passive gleaning).

The effects of clutter on biosonar performance are reflected in a number of behavioural studies in bats: Warnecke et al. (2014) showed that target detection was impaired when target and clutter were arranged along the same azimuth and elevation, but shifting the clutter source off-axis lead to a spatial release from masking and facilitated target detection. Moss et al. (2011) showed how free-flying bats negotiate such spatial unmasking and temporal resolution to locate and intercept prey in a complex environment. These behavioural experiments provided spatial cues not only along the distance axis, but also along the azimuth and/or elevation axes (bats could adjust their flight paths). However, backward- and forward masking of the clutter onto the suspended target - and thus the bats’ capability to perceptually resolve target and clutter along the distance axis - will contribute to the bats’ performance.

Recently, Geipel et al. (2019) investigated a bat species that hunts silent and motionless prey among dense vegetation, solely using echolocation. The authors tested a hypothesis (modified after Denzinger and Schnitzler (2013)) stating that such a foraging strategy would exploit “an isolated additional [prey] echo between the clutter echoes” (Denzinger and Schnitzler, 2013). This describes exactly the current results, where bats had to detect the target reflection between the masker reflections. As has been pointed out earlier though (Baier, 2019), the study by Geipel et al. (2019) found maximum delays of 0.25 ms between the target and the clutter echoes, since prey items were perched directly on leaves. Considering the current results, it would be impossible for these bats to separate the echoes in the time domain. Accordingly, the authors investigated and confirmed a different strategy, namely the active reduction of clutter echoes by approaching from angles that transform the leaf into a specular reflector (Geipel et al., 2019).

### Acoustic properties of echolocation signals

Biosonar is an active sense, i.e. the animal itself produces the signal with which it probes its environment. Like all microchiropteran bats, *Phyllostomid* bats produce their echolocation calls with exceptionally thin vocal folds and use their vocal tract to further shift energy into ultrasonic frequency ranges, often producing multiharmonic calls.

Within species-specific limits, bats can change the temporal and spectral properties of their signals according to the perceptual task at hand. Temporal parameters like call duration concern call timing, spectral parameters like minimum and maximum frequency concern call frequency-content. Generally speaking, bats produce shorter and broader/higher calls in cluttered environments. Shorter call durations result in more accurate and up-to-date information due to less overlap between consecutive echoes (or call and echo!) even at high repetition rates. Bats tend to avoid overlap of target echo and clutter echoes as well as overlap of call and echo (Kalko and Schnitzler, 1993; Kalko and Schnitzler, 1989). Higher call frequencies result in higher spatial acuity due to shorter wavelengths and higher directionality (Griffin, 1958). The range of frequencies that an echolocation call covers, its bandwidth, determines accuracy in ranging (Simmons, 1973). Within the genus of *Myotis* bats, those species with the largest bandwidth are most successful to find prey suspended in front of a clutter surface (Siemers and Schnitzler, 2004).

In light of this, we analysed the echolocation calls that the bats used throughout the experiment. We found no evidence that bats adapted their call parameters in response to task difficulty or in relation to their individual resolution limit (Fig. 5).

However, the call durations we observed are in line with values observed in *P. discolor* bats when echolocating towards an approaching food source (Linnenschmidt and Wiegrebe, 2016). When echolocating toward the target at 100–110 cm distance, bats used call durations of 0.3–0.9 ms (cf. Fig.2 in Linnenschmidt and Wiegrebe (2016). The call durations we found in the current experiment (around 0.4-0.7 ms) are a good match given a simulated distance of 107 cm between the virtual target and the bat. Due to the difference in target and background structure between the studies, we do not compare spectral parameters of the current study to those in Linnenschmidt and Wiegrebe (2016).

In summary, our work offers compelling evidence for spatial resolution along the distance axis in echolocating bats. Corroborating earlier work with a more robust experimental design, we have introduced a virtual-reality approach with complex echo-acoustic scenarios, precluding non-conclusive perceptual cues. We demonstrated that *Phyllostomus discolor* bats can listen into a perceptual dip between multiple reflections and therefore possess range resolution.

## List of abbreviations

2AFC: two-alternative, forced choice
FM: frequency-modulated
IR: impulse response
IMD: inter-masker delay (onset difference between 1^st^ and 2^nd^ masker reflection)
SC: spectral centroid (weighted mean of frequencies present in the signal)

## Acknowledgements

For help during data acquisition, we thank E. Lattenkamp and E. Mardus. For help with analysis in R, we thank F. Korner-Nievergelt. We are grateful to B. Grothe for providing excellent research infrastructure. All experiments complied with the principles of laboratory animal care and were conducted under the regulations of the current version of the German Law on Animal Protection (approval 55.2-1-54-2532-34-2015, Regierung von Oberbayern). This study is dedicated to the memory of our co-author, mentor and friend Lutz Wiegrebe. His ceaseless curiosity and selfless support will remain a perpetual inspiration.

## Author contributions

Conceptualization, LW; Methodology, LW and ALB; Software, LW and ALB; Formal Analysis, PAW, LW and ALB; Investigation, PAW and ALB; Writing – Original Draft, LW and ALB; Writing – Review & Editing, ALB; Visualization, PAW and ALB; Supervision, LW and ALB; Project Administration, LW and ALB; Funding Acquisition, LW.

## Data availability

Data will be made publicly available at the time of publication.

